# Temporal transcriptomic changes during neurodevelopment in a mouse model of Smith-Lemli-Opitz syndrome

**DOI:** 10.1101/2023.11.21.568116

**Authors:** Amy Li, Hideaki Tomita, Libin Xu

## Abstract

Smith-Lemli-Opitz syndrome (SLOS) is a cholesterol biosynthesis disorder that arises from mutations in the gene *DHCR7*, leading to decreased production of cholesterol and accumulation of its precursor, 7-dehydrocholesterol. SLOS displays a wide range of neurodevelopmental defects, intellectual disability, and behavioral problems. However, an in-depth study on the temporal changes of gene expression in the developing brains has not been done before. In this work, we carried out the transcriptomic analysis of whole brains from WT and *Dhcr7*-KO mice at embryonic day 12.5 (E12.5), E14.5, E16.5, and postnatal day 0 (PND0). First, we observed the expected downregulation of the *Dhcr7* gene in the *Dhcr7*-KO brains, as well as changes of other genes involved in cholesterol biosynthesis at all time points. Pathway and GO term enrichment analyses revealed affected signaling pathways and biological processes that were shared amongst time points and unique to individual time points. Specifically, the pathways important for embryonic and neural development, including Hippo, Wnt, and TGF-β signaling pathways, are the most significantly affected at the earliest time point, E12.5. Additionally, neurogenesis-related GO terms were enriched in earlier time points, consistent with the timing of development. Conversely, pathways related to synaptogenesis, which occurs later in development compared to neurogenesis, are significantly affected at the later time points, E16.5 and PND0, including the cholinergic, glutamatergic, and GABAergic synapses. *In vitro* neurogenesis experiments using GABAergic neuronal precursors isolated from embryonic mouse brain confirmed that loss of *Dhcr7* led to decreased proliferation and premature neurogenesis, consistent with the transcriptomic changes.

## 1. Introduction

Smith-Lemli-Opitz syndrome (SLOS, OMIM #270400) is caused by genetic mutations in the *DHCR7* gene [1–5], which encodes the terminal enzyme of cholesterol biosynthesis, 7-dehydrocholesterol reductase (DHCR7, EC 1.3.1.21), leading to decreased levels of cholesterol and increased levels of the cholesterol precursor, 7-dehydrocholesterol (7-DHC), other sterol intermediates, and oxysterol metabolites [6–8]. Clinically, the disorder manifests a wide variety of phenotypes, including multiple congenital malformations (structural abnormalities and functional defects across several organ systems), developmental delay, cognitive impairment, and autistic behavior [9–11]. Specifically in the CNS, anatomical abnormalities have been observed in SLOS patients, ranging from microcephaly to enlarged ventricles and hypoplasia of the corpus callosum, and in the most severe cases, holoprosencephaly [6, 12–14].

Development of the CNS is a carefully orchestrated and regulated process involving the interplay of many different signaling pathways. Within the CNS, cholesterol plays an essential role as a precursor to neurosteroids and steroid hormones, a major component of plasma membranes, myelin, and lipid rafts, and an important modulator of developmental signaling pathways, such as the Sonic Hedgehog pathway [15–18]. Thus, disruption of cholesterol synthesis profoundly impacts many cellular processes in brain development, given the numerous biological functions of cholesterol. In addition, the cholesterol pool in the brain is especially unique due to the blood-brain barrier preventing cholesterol uptake from the circulation [19]. This means that cholesterol metabolism must be tightly regulated in a spatiotemporal manner during neurodevelopment to supply the high metabolic needs of cells during neurogenesis, as cells proliferate, migrate, differentiate, and undergo programmed cell death [19, 20]. While there is limited maternal-fetal transfer during early development in mice, *de novo* cholesterol synthesis becomes critical in the CNS after the formation of the blood-brain barrier around E10-11 [21], while cortical neurogenesis begins around E12.5 and peaks around E15.5.

The development of genetically engineered mouse models of SLOS has made it possible to study the molecular mechanisms underlying the disorder at embryonic stages. In this study, we utilize a commercially available SLOS mouse model, *Dhcr7*^-/-^ or *Dhcr7*-KO (Ex8), in which the exon VIII coding region of *Dhcr7* has been deleted, leading to a truncated gene product [22]. The mouse model replicates the known biochemical defect of cholesterol synthesis [23] and exhibits developmental abnormalities in the lung, cleft palate, and bladder [22]. Homozygous pups die on the first day of birth due to an uncoordinated suck and failure to feed.

Transcriptomic studies of SLOS models have been previously reported in the literature [23–25]. Waage-Baudet et al. carried out the first transcriptomic study of the hindbrains of the same *Dhcr7*-KO SLOS mouse model, but on the 129/SvEv background, at gestational day 14 (E14) using the Affymetrix microarray analysis [24]. Hierarchical clustering of the differentially expressed genes identified alterations in genes involved in cholesterol homeostasis, cell cycle control and apoptosis, neurodifferentiation, embryogenesis, and, of particular interest, axon guidance. Dysregulated axon guidance was believed to be partly responsible for the abnormal hippocampal development that was previously reported in these mice [26]. In another study using a human-derived induced pluripotent stem cell (iPSC) model of SLOS that exhibited aberrant neural differentiation, Francis et al. determined that several transcriptional networks related to neural differentiation, cadherin-associated signaling, and kinase signaling were affected when comparing SLOS iPSCs to control iPSCs. In addition, they found overall downregulation of Wnt/β-catenin signaling (*CAV1, CDH1, SNAI2*), suggesting a role of defective Wnt signaling in the aberrant iPSC differentiation phenotype in SLOS [25].

In a recent study from our lab, we found that *Dhcr7*-KO leads to a premature neurogenesis (with decreased proliferation and increased differentiation) of cortical neural precursors in both a SLOS mouse model and in SLOS patient-derived neural progenitor cells (NPCs) and eventual thinning of cortical layers in embryonic mouse brains [23]. This mechanism was found to be mediated by glucocorticoid receptor and the neurotrophin kinase, TrkB. RNA sequencing analysis of human SLOS NPCs revealed gene expression changes related to neural precursor proliferation and differentiation, including those involved in MAPK and Ras signaling. While some transcriptomics experiments have been performed in SLOS models, no study has examined the temporal changes in gene expression and biological pathways throughout the embryonic developmental stages.

In this work, we carried out transcriptomic analysis of whole brains [27] in a well-established SLOS mouse model, *Dhcr7*-KO mice, at E12.5, E14.5, E16.5, and PND0 in comparison with matching wild-type (WT) mice. Differentially expressed genes and significantly affected signaling pathways and biological processes were identified at each time point. Meta-analysis of all four time points highlighted shared and unique biological pathways at distinct developmental stages. *In vitro* experiments validated some of the transcriptomic findings, such as decreased proliferation and increased neurogenesis at the early time points, E12.5 and E14.5.

## 2. Methods and Materials

### 2.1 Chemicals and biochemicals

*2-methylbutane was purchased from Thermo Fisher Scientific (Grand Island, New York). RNeasy Lipid Tissue Mini Kit was purchased from Qiagen (Germantown, Maryland)*. The primary antibodies used for immunostaining were chicken anti-GFP (1:1000; Abcam), rabbit anti-Dhcr7 (1:100; Thermo Fisher Scientific), mouse anti-Ki67 (1:200; BD Biosciences), mouse anti-βIII-tubulin (1:1000; Covance), rabbit anti-βIII-tubulin (1:1000; Covance), rabbit anti-cleaved caspase 3 (1:400; Cell Signaling Technology). The secondary antibodies used for immunostaining were Rhodamine (TRITC)-conjugated goat anti-mouse and anti-rabbit IgG (1:500; Jackson ImmunoResearch Laboratories) and Alexa Fluor 488-conjugated goat anti-mouse and anti-rabbit IgG (1:800; Jackson ImmunoResearch Laboratories).

### 2.2 Animals

Animal studies were performed in accordance with the NIH *Guide for the Care and Use of Laboratory Animals* and approved by the University of Washington Institutional Animal Care and Use Committee. C57BL/6J and transgenic heterozygous mice with a null mutation for *Dhcr7* (Ex8) mice were purchased from Jackson Laboratories (Bar Harbor, Maine; catalog #007453). The genotype primers used are the following: *Dhcr7*-WT-F: 5’-GGATCTTCTGAGGGCAGCTT-3’; *Dhcr7*-WT-R: 5’-TCTGAACCCTTGGCTGATC-3’; Delta-Mut: 5’-CTAGACCGCGGCTAGAGAAT-3’. Mice were housed in an animal care facility with a standard 12-hour light and dark cycle and fed an *ad libitum* commercial rodent chow diet. Heterozygous *Dhcr7* (Ex8) mice were mated overnight, where the day after time-mating was designated embryonic day 0.5 (E0.5). Heterozygous mating produced *Dhcr7*^+/+^ (WT), *Dhcr7*^+/-^ (Het), and *Dhcr7*^-/-^ (KO) offspring at roughly the expected Mendelian 1:2:1 ratio. Animals were genotyped using PCR as described previously [22]. Brains were harvested from mice at E12.5, E14.5, E16.5 (n=3 per genotype), and PND0 (n=4 per genotype) time points. After euthanasia, whole brains were excised under the dissection scope in ice-cold PBS and frozen in pre-cooled 2-methylbutane to be stored at -80°C until further processing steps.

### 2.3 RNA Isolation

E12.5, E14.5, and E16.5 whole brains and PND0 half brains (cut along the midsagittal plane) were processed for RNA sequencing. RNA was isolated using the Qiagen Lipid RNeasy Kit following the manufacturer’s protocol. RNA quality was assessed with a NanoDrop One spectrophotometer (Thermo Scientific), and only samples with appropriate purity (260/280 ratio > 2.0 and 260/230 ratio > 2.0) were used for analysis. Gel electrophoresis was also used to check for DNA contamination and RNA integrity.

### 2.4 RNA Sequencing (Novogene)

Samples were sent to Novogene Co. Ltd (Sacramento, CA) for total RNA sequencing on their Illumina platform. RNA sample quality control was further assessed, and the mRNA library was prepared with additional polyA enrichment, RNA fragmentation, and cDNA transcription steps. Libraries were sequenced with 150 bp paired-end reads on the Illumina NovaSeq 6000 Sequencing System with a minimum sequencing read depth of 6 Gb of raw data per sample.

### 2.5 RNA Sequencing Data Analysis

Data analysis was performed using a combination of Linux, R code, and various open-source online tools. We used an in-house-built Python script that streamlines raw data processing steps, utilizing zipped raw sequence read files to conserve disk memory space. *hisat2* is used to align sequence reads to the reference mouse genome (mm10), where greater than 95% sequence alignment was achieved [28]. *samtools sort* and *featureCounts* are then used to count reads to genomic features (annotation file VM25) [29, 30]. The resulting sequence count data are input into DESeq2 (using R/Bioconductor) for differential expression analysis [31]. DESeq2 uses a generalized linear model with negative binomial distribution to estimate dispersion and the Wald significance test. Independent filtering step removes genes with very low counts. For pathway analysis, we utilized iPathwayGuide and Database for Annotation, Visualization, and Interpretation (DAVID) [32–35]. For both iPathwayGuide and DAVID, we used a list of all measured genes specific for each time point as the reference background lists.

### 2.6 Cortical precursor cell cultures

Mouse neural precursor cells were isolated from the medial ganglionic eminence (MGE) and the lateral ganglionic eminence (LGE) and were cultured as described previously [36, 37]. Briefly, the MGE/LGE was dissected from E11.5 WT or *Dhcr7*^-/-^ (KO) mouse embryos in ice-cold HBSS (Invitrogen) and transferred into cortical precursor medium. The neural precursor medium consisted of Neurobasal medium (Invitrogen) with 500 µM L-glutamine (Invitrogen), 2% B27 supplement (Invitrogen), 1% penicillin-streptomycin (Invitrogen), and 40 ng/ml fibroblast growth factor 2 (FGF2) (Sigma). The dissected MGE/LGE was mechanically dissociated by a fire-polished glass pipette and plated onto 24-well plates coated with 2% laminin (BD Biosciences) and 1% poly-D-lysine (Sigma-Aldrich). Plating density of the cortical precursors was 150,000 cells/well in 24-well plates for single embryo cultures.

### 2.7 Neurosphere cultures

E11.5 MGE/LGE from WT or *Dhcr7*^-/-^ embryos were isolated and mechanically dissociated into a single cell suspension by fire-polished glass pipette as previously described [37, 38]. Cell density and viability were determined using trypan blue exclusion. Cells were seeded in triplicate at a clonal density of 10 cells/μl in 6 well (2 ml/well) ultra-low attachment culture plates (Coster) in serum-free medium supplemented with 20 ng/ml epidermal Growth Factor (EGF) (Sigma), 20 ng/ml FGF2 (Sigma), 2% B27 supplement (Invitrogen) and 2 μg/ml heparin (Sigma). Neurospheres were cultured for 6 days at 37 °C. To evaluate self-renewal potential, neurospheres were mechanically dissociated into single cell suspensions by fire-polished glass pipette, passed through a 45-μm nylon screen cell strainer, and cultured at a clonal density of 2 cells/μl for an additional 6 days.

### 2.8 Oxysterol and Sterol Analysis

Cell pellets were resuspended in 300 μL of 1X PBS and lysed by sonication in an ice bath for 30 min. Protein determination was performed using the BioRad DC protein mass assay (BioRad, Hercules, CA). Internal standard mixtures for sterols and oxysterols analysis were added to each sample (see [39]). Lipid extraction was performed using the Folch method as described previously [40–42]. The dried lipid extract was reconstituted with 200 μL of methylene chloride and stored at -80 °C until analysis. Prior to analysis, 50 μL of extract was transferred into glass LC vials, dried under argon, and reconstituted with 50 μL of 90% methanol in water with 0.1% formic acid. Determination of oxysterol and sterol concentrations in cells and tissues was performed by ultra-performance liquid chromatography (UPLC) tandem mass spectrometry (MS/MS) on a SCIEX 6500 triple quadrupole mass spectrometer (for oxysterols) or a SCIEX 4000 QTRAP (for sterols) mass spectrometer with atmospheric pressure chemical ionization (APCI) coupled to a Waters Acquity UPLC system, as described previously [41, 42]. Briefly, sterols and oxysterols were separated by reverse-phase chromatography on a C18 column (1.7 mm, 2.1 x 100 mm, Phenomenex Kinetex) using a 15 min isocratic gradient of 90% methanol with 0.1% formic acid at a flow of 0.4 mL/min. Selective reaction monitoring (SRM) was used to monitor the dehydration of the sterol and oxysterol [M+H]^+^ ions to generate [M+H-H_2_O]^+^ ions [39]. The APCI parameters were as follows: nebulizer current, 3 mA; temperature, 350 °C; curtain gas, 20 psi; ion source gas, 55 psi. The MS conditions for SRM analysis were as follows: declustering potential, 80 V; entrance potential, 10 V; collision energy, 25 V; collision cell exit potential, 20 V. Data analysis was performed with Analyst (v. 1.6.2) Quantitation Wizard as described previously [39]. Concentrations were normalized to cell protein weight or tissue weight.

### 2.9 Data Availability

Raw RNA-seq data have been deposited in NCBI’s Gene Expression Omnibus (GEO) and are available with the GEO Accession Number: GSE247566.

## 3. Results

### 3.1 Differential Expression Analysis

RNA sequencing (RNAseq) of whole mouse brains was performed, and gene expression profiles were compared between WT and *Dhcr7*-KO mice at three embryonic time points (E12.5, E14.5, E16.5) and one postnatal time point (PND0). RNAseq steps are outlined in **Figure 1**, including brain dissections, RNA extraction and sequencing, and data analysis.

**Figure 1.**
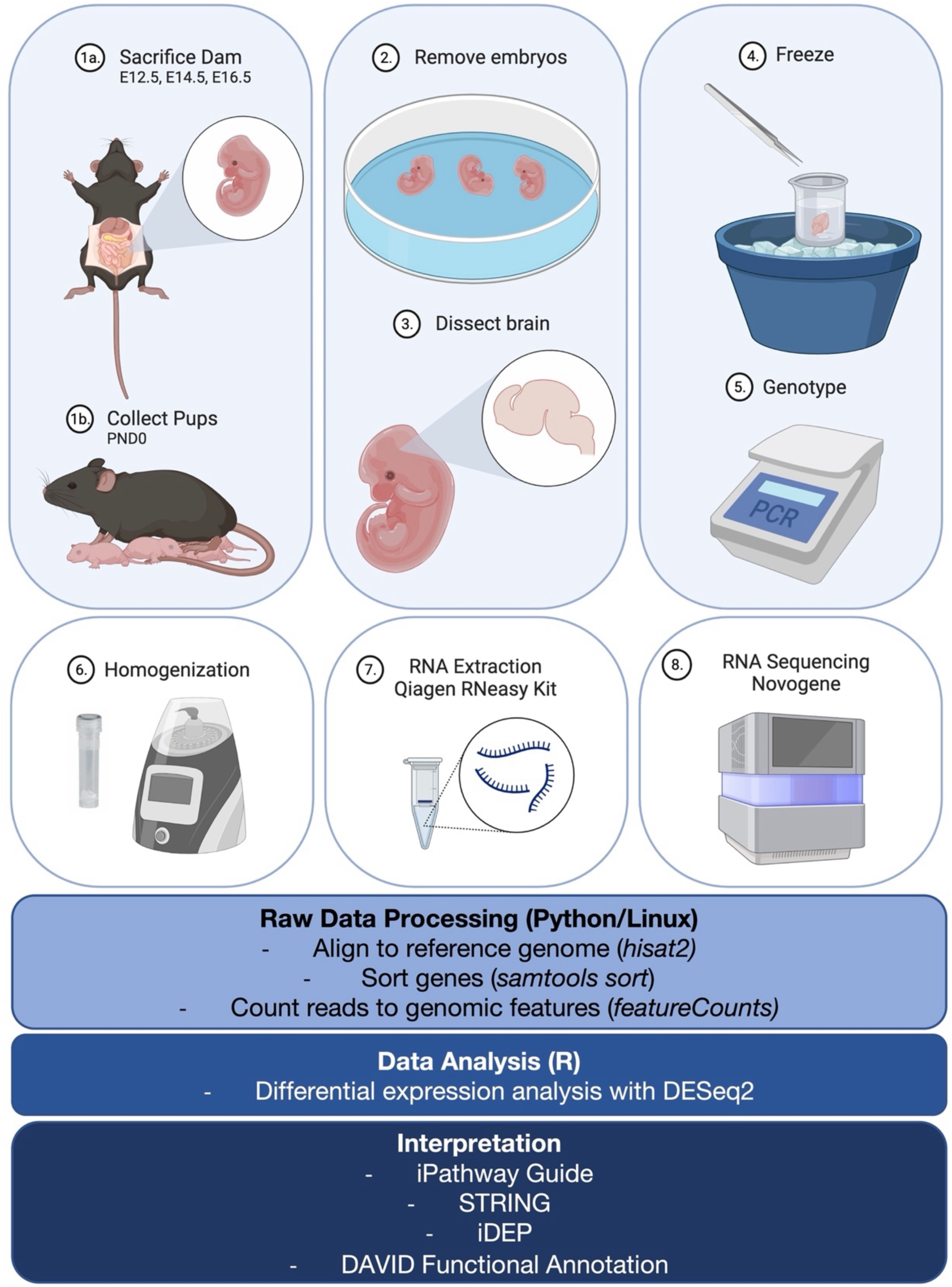
Diagram of RNA sequencing workflow. Steps for sample preparation (brain tissue collection from mouse embryos or neonates), RNA extraction, RNA sequencing, and data analysis are outlined. Created with BioRender.com.

When considering all the WT and KO samples together in one principal component analysis (PCA) bi-plot (**Supplementary Figure S1A)**, there is separation and clustering of the four groups according to time points, indicating that there are differences in the transcriptome at each time point that make each cluster distinct from the others. The PCA bi-plots constructed for each individual time point (**Supplementary Figure S1B**) more clearly demonstrate the separation between WT and KO samples.

To examine gene expression differences between the WT and KO conditions at the various time points, we performed differential expression analysis using DESeq2. The number of differentially expressed genes (DEGs) for each time point, using an adjusted p-value (or false discovery rate, FDR) threshold of 0.05, are summarized in **Table 1**, including the number of total measured genes used as a reference background for pathway analysis. In summary, E12.5 had 647 differentially expressed genes (213 up-regulated in *Dhcr7*-KO compared to control, 434 down-regulated), E14.5 had 279 genes (129 up-regulated, 150 down-regulated), E16.5 had 645 genes (391 up-regulated, 254 down-regulated), and PND0 had 919 genes (391 up-regulated, 528 down-regulated). In **Figure 2**, significant DEGs (using thresholds: adjusted p-value (FDR) ≤ 0.05 and absolute fold change ≥ 1.2) are visually represented on volcano plots displaying the magnitude of change on the y-axis and statistical significance on the x-axis. The complete list of DEGs is available in **Supplementary Table S1**. The low number of measured genes for E14.5 resulted from the independent filtering step in DESeq2, which removes features with very low read counts that have low power to be detected as significant due to high dispersion.

**Figure 2.**
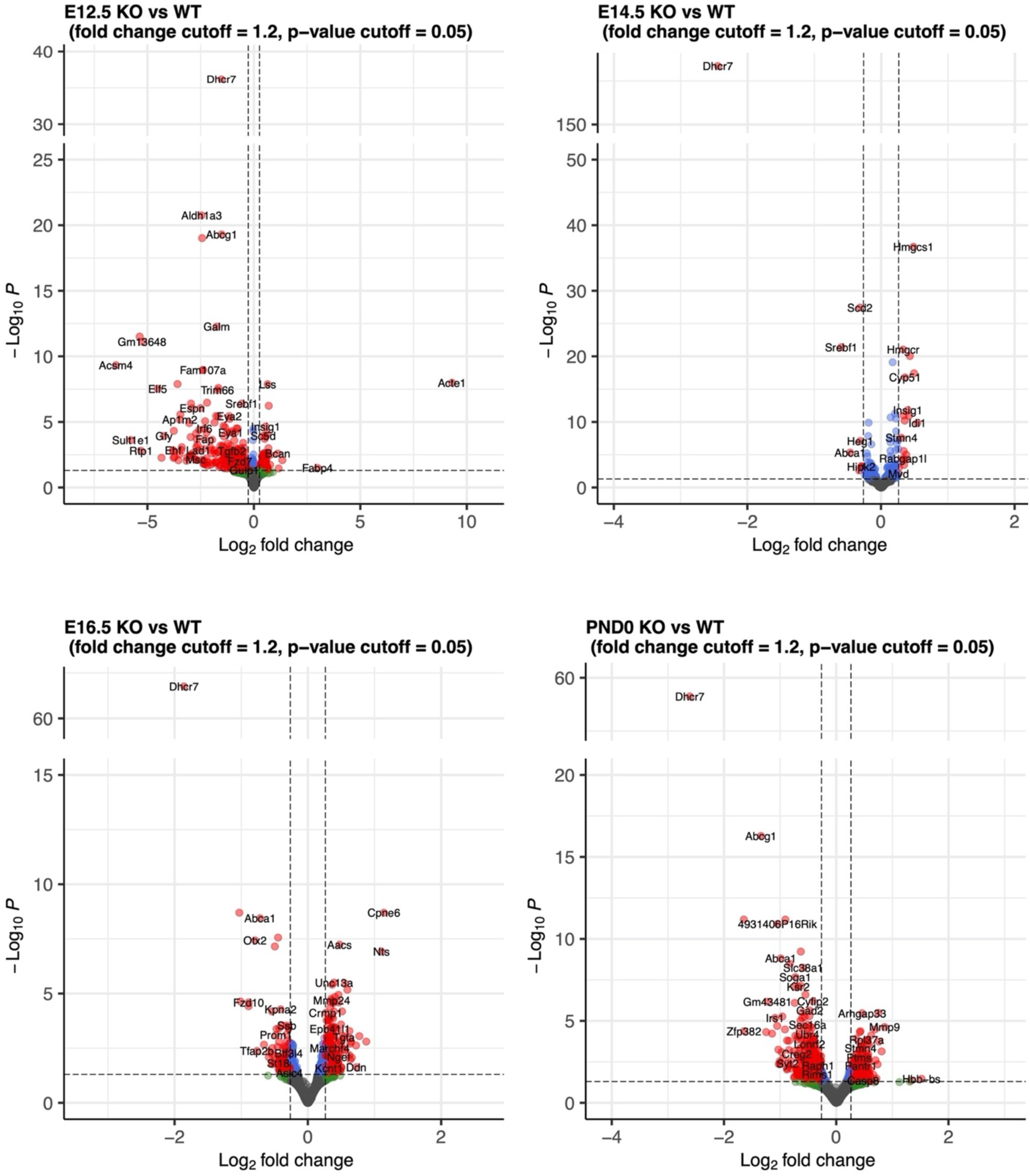
Volcano plots highlighting differentially expressed genes (shown in red) filtered by adjusted p-value ≤0.05 (shown in blue) and fold change ≥1.2 (shown in green). The complete list of DEGs is available in **Supplementary Table S1**. Created with R/*Enhanced Volcano*.

**Table 1.** Summary of differentially expressed genes, with an adjusted p-value threshold (FDR ≤ 0.05), or an adjusted p-value threshold and a fold change threshold (|FC| ≥ 1.2), and the total number of genes with measured expression (after DESeq2 filtering of genes with an extreme count outlier and low mean normalized counts).

| Time Point | Number of DEGs<br>($FDR \leq 0.05$) | Number of DEGs<br>($FDR \leq 0.05$ and $\vert FC\vert \geq 1.2$) | Total Number<br>of Measured<br>Genes |
| --- | --- | --- | --- |
| E12.5 | 647 (Up: 213; Down: 434) | 644 (Up: 212; Down: 432) | 17386 |
| E14.5 | 279 (Up: 129; Down: 150) | 42 (Up: 29; Down: 13) | 6559 |
| E16.5 | 645 (Up: 391; Down: 254) | 470 (Up: 297, Down: 173) | 11674 |
| PND0 | 919 (Up: 391; Down: 528) | 844 (Up: 369; Down: 515) | 13145 |

Interestingly, most DEGs did not overlap between time points, as seen in **Figure 3**, suggesting that there are differences in the genes and pathways being affected at specific times throughout development. Only nine DEGs were found to be shared between all four groups. Three that were down-regulated (*Abca1, Dhcr7, Srebf1*) and six up-regulated (*Aacs, Fdft1, Hmgcs1, Nsdhl, Sqle, Stmn4*). Except for *Stmn4*, the rest of the genes are all directly involved in different aspects of sterol metabolism, such as efflux transport (*Abca1*), transcriptional regulation (*Srebf1*), and various enzymes along the synthetic pathway. As expected in this *Dhcr7*-KO mouse model, the gene encoding *Dhcr7* was the most down-regulated gene transcript, with the largest fold changes across all four time points and the lowest *p*-values. However, the other observed changes, such as up-regulation of key enzymes involved in cholesterol biosynthesis, are likely compensatory mechanisms that are transcriptionally regulated by the low cellular cholesterol levels in *Dhcr7*-KO mice.

**Figure 3.**
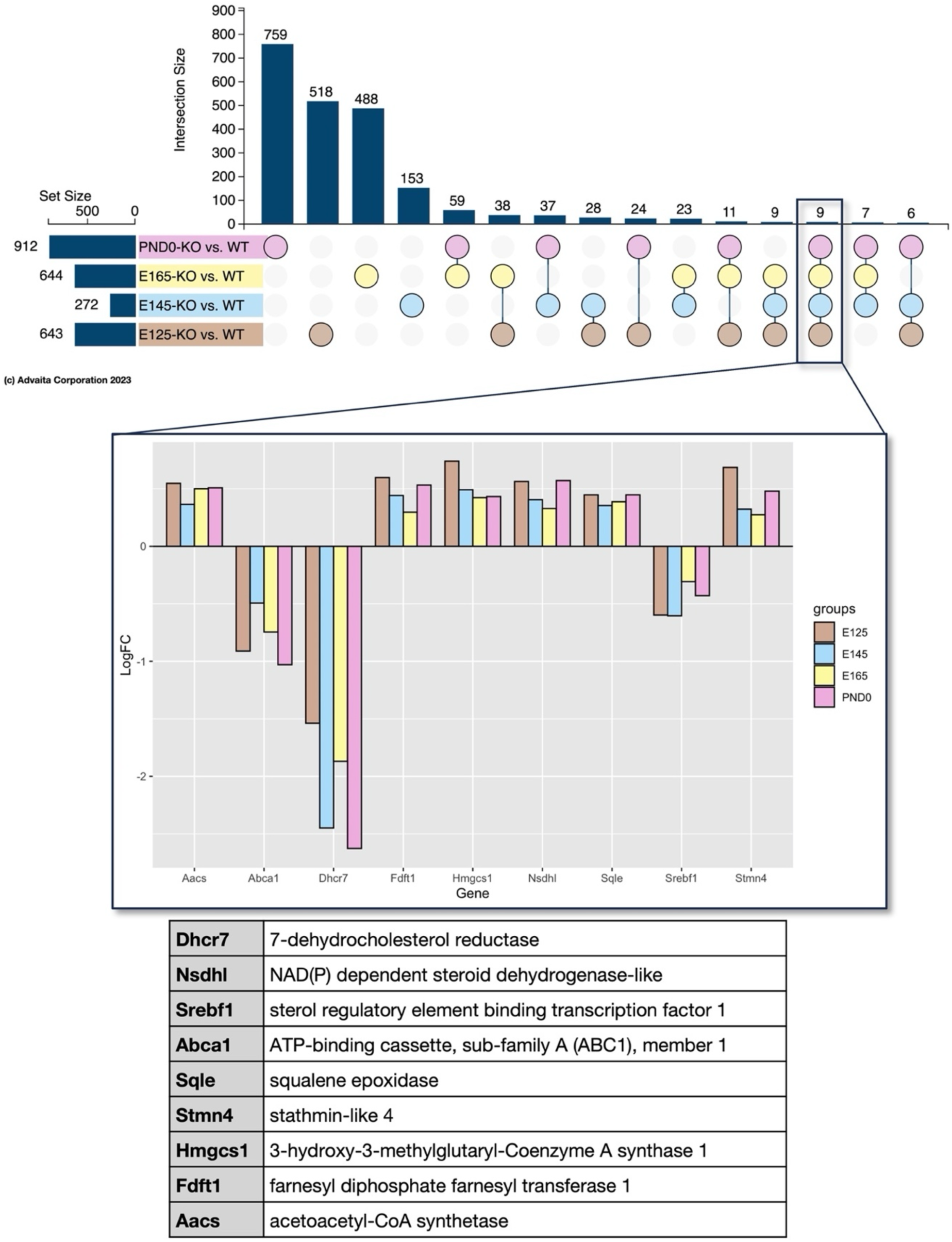
UpSet plot for meta-analysis of differentially expressed genes from all four time points. Bar plot displaying log fold-change values for the nine shared genes from all four time points and table of gene annotations. Created by iPathwayGuide and R/*ggplot2*.

### 3.2 Pathway Analysis and Gene Ontology (GO) Term Enrichment

For genes included in pathway analysis, we used a significance threshold of an adjusted *p*-value (FDR) ≤ 0.05. Pathway analysis is used to identify which pathways are over-represented in a given set of genes beyond pure random chance. iPathwayGuide uses its proprietary “Impact Analysis” that factors in two probabilities, pORA (over-representation) and pAcc (perturbation) to compute statistical significance and is modelled after the Kyoto Encyclopedia of Genes and Genomes (KEGG) pathways [34, 35, 43]. Many pathways were significantly affected at individual time points. The top 10 KEGG pathways significantly affected at each time point are shown in **Figure 4**, and additional significantly altered pathways are available in **Supplementary Table S2.** Pathway results are visualized with bubble plots, which are ranked by *p*-value. The horizontal position on the plot designates the ratio of differentially expressed genes to the total number of genes associated with the specific pathway, while the size of the bubble designates the number of differentially expressed genes and the color designates the *p*-value. Interestingly, some of the top results in the earliest time point (E12.5) include important signaling pathways for embryonic development (Hippo, Wnt, and TGF-β). In contrast, later time points include pathways related to neurotransmission (synaptic vesicle cycle and different types of synapses, including cholinergic, glutamatergic, and GABAergic synapses).

**Figure 4.**
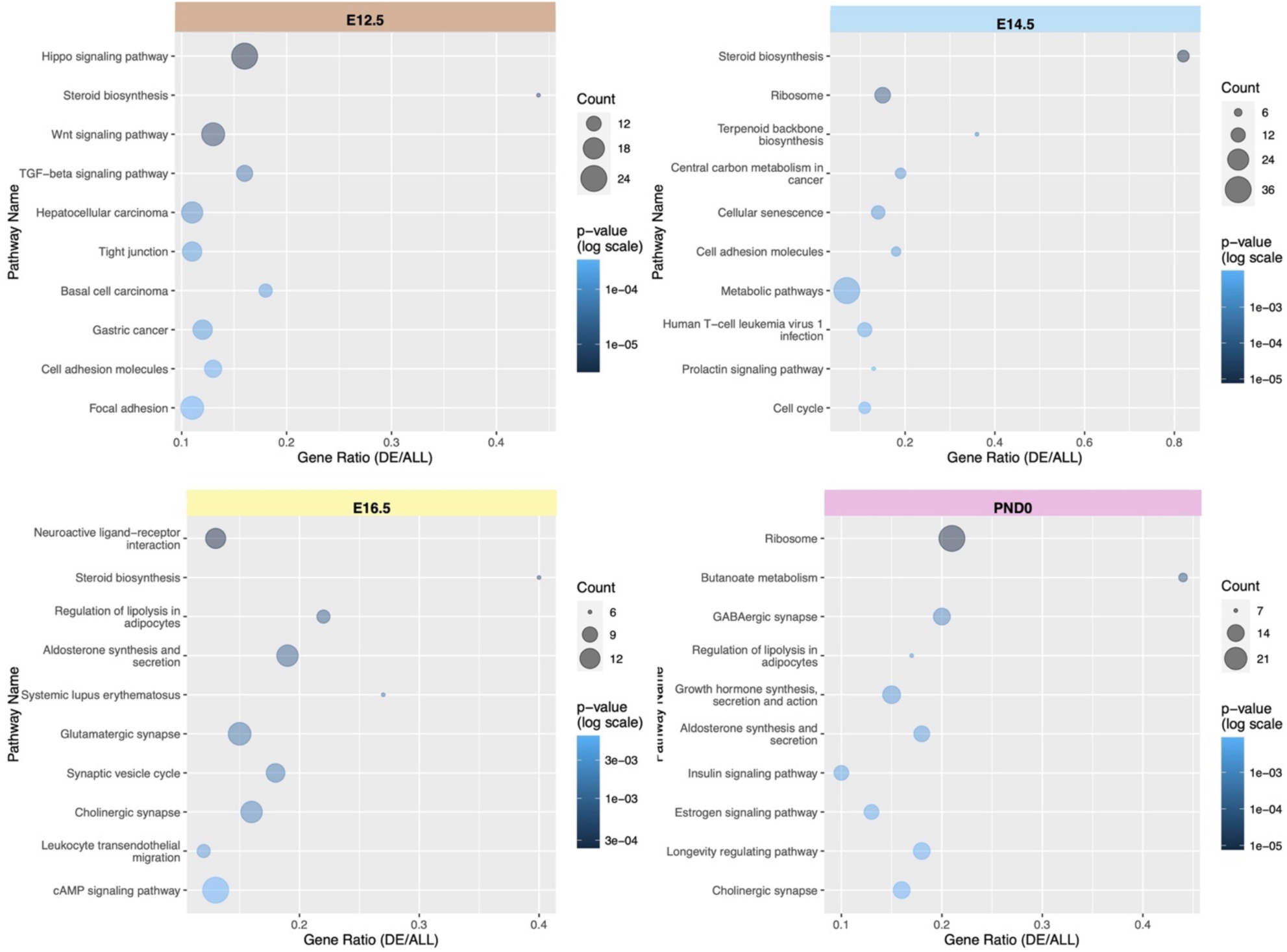
Top 10 enriched pathways from pathway analysis using iPathwayGuide. Bubble plots are ordered by p-value, and display pathway name and gene ratio, where the size of the bubble designates the number of differentially expressed genes, and the color designates the p-value. * COVID-19 removed from E14.5, E16.5, and PND0 time points. Created by R/ggplot2.

We also performed a GO enrichment analysis to identify which GO terms for biological processes are significantly over-represented in the given set of genes [44, 45]. The top 10 GO biological process terms enriched at each time point are shown in **Supplementary Figure S2**. Like pathway analysis, we found many of the top terms related to sterol synthesis, development, and synaptic signaling. The complete list of significantly enriched GO terms for biological processes is available in **Supplementary Table S3.**

### 3.3 Pathway Meta-Analysis of All Time Points

The meta-analysis feature in iPathwayGuide enables cross-comparison of the four time points, as shown via the UpSet plot in **Figure 5**. *Steroid biosynthesis* (mmu:00100) is the only pathway significantly affected at all time points, with a p-value significance threshold set at 0.05. This is a similarly expected result as the shared DEGs. **Figure 6** shows the genes associated with the *steroid biosynthesis pathway* and their log fold-changes for each time point. This list of genes includes transcripts different from the nine shared DEGs (from **Figure 3**), including *Lss, Msmo1, Cyp51, Sc5d, Dhcr24, and Tm7sf2.* In this case, *Dhcr7* is the only down-regulated gene, while all the other steroid synthesis-related genes are up-regulated, indicating compensatory response.

**Figure 5.**
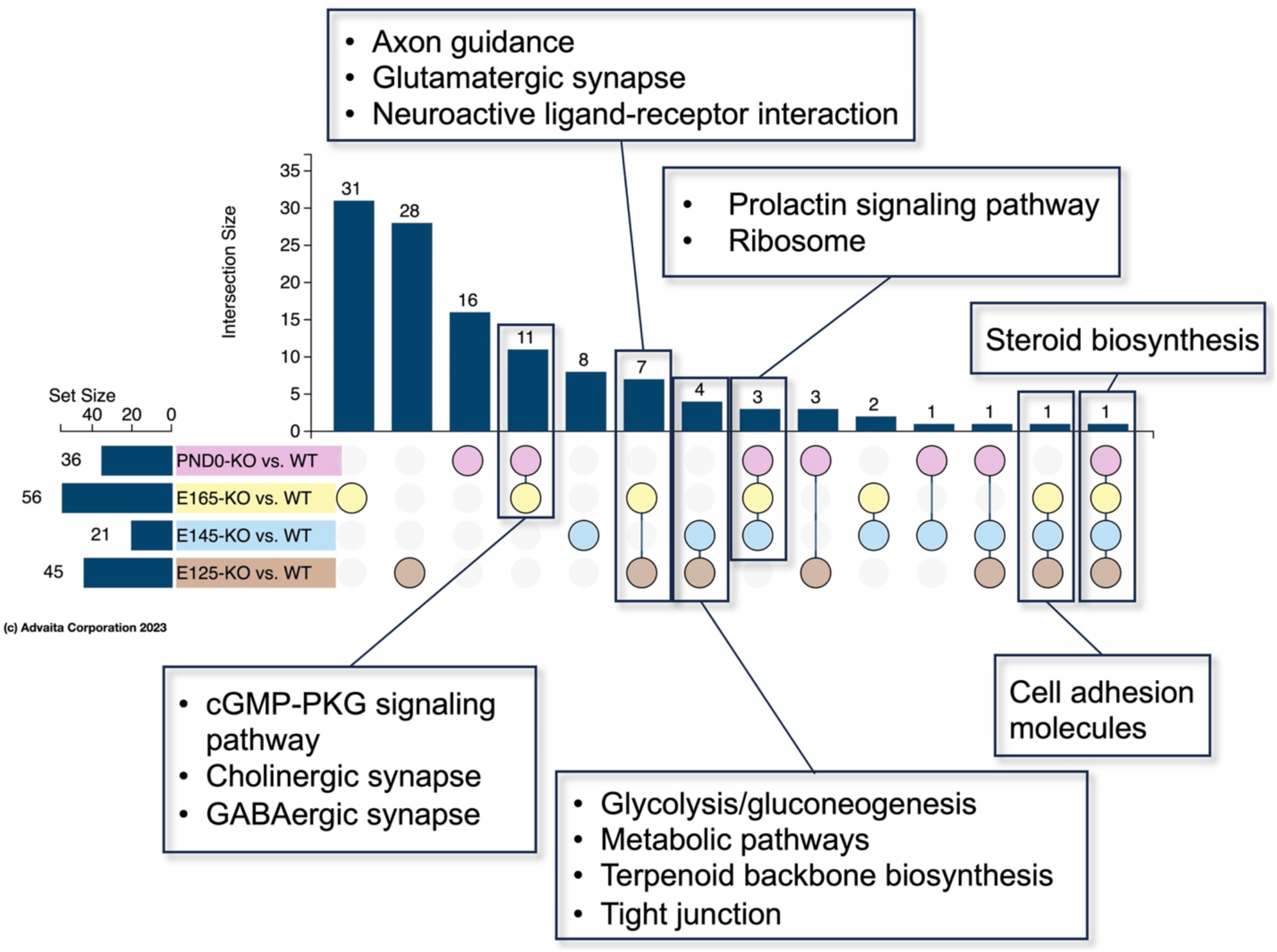
UpSet plot for meta-analysis of shared pathway hits from all time points with some specific pathways listed. All pathway results are available in **Supplementary Table S2**. Created by iPathwayGuide.

**Figure 6.**
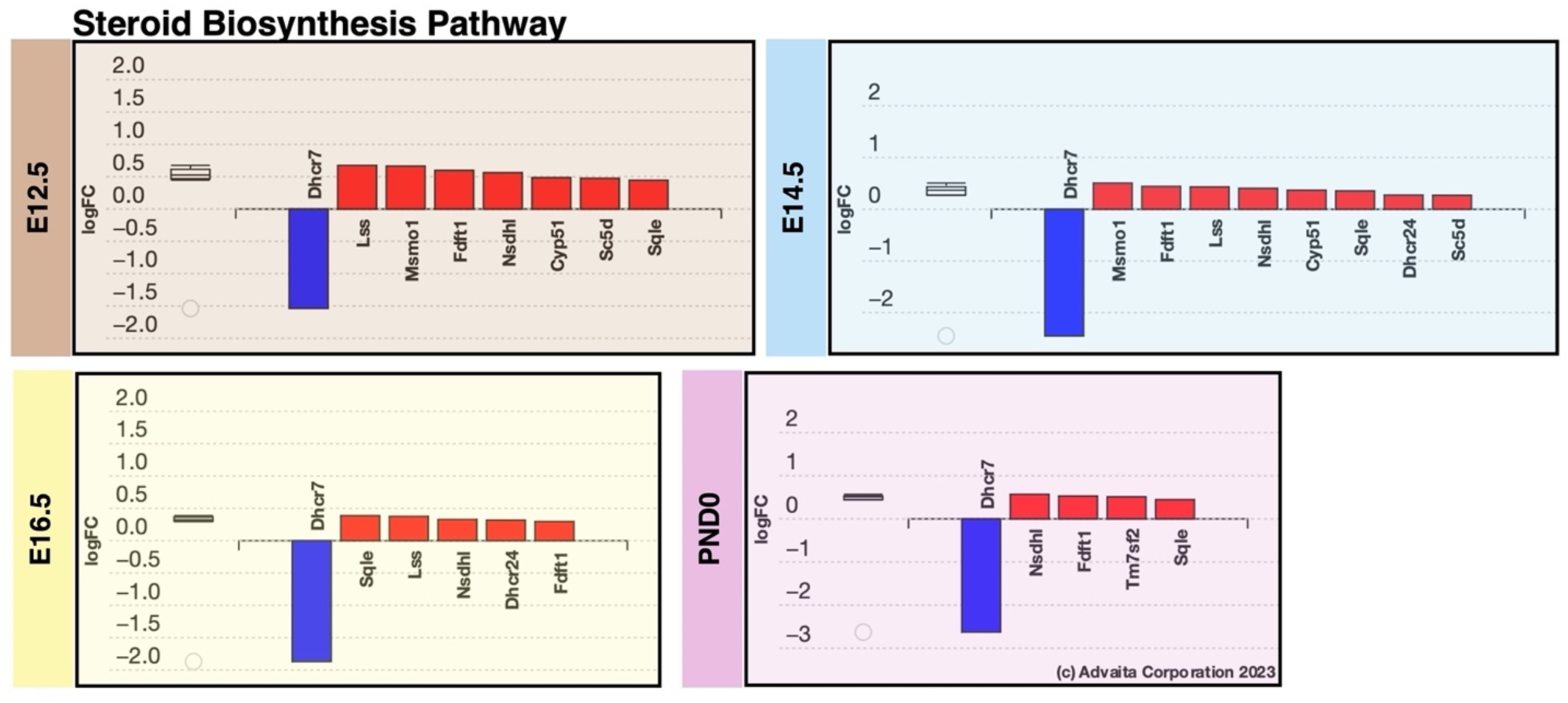
Bar plots of log fold-changes for DEGs in the **steroid biosynthesis pathway** (mmu:00100) for time points where this pathway had a p-value ≤ 0.05. DEGs are ordered by absolute LFC.

In **Figure 5**, we have highlighted pathways that are commonly affected at two or more time points. For example, the *cell adhesion molecules pathway* (mmu:04515) is the only other pathway shared between all embryonic time points, E12.5-E16.5 (**Figure 7**). *L1cam* is the only gene consistently up-regulated at those three time points. The *ribosome pathway* (mmu:03010) was significantly affected in E14.5, E16.5, and PND0 groups, and the associated genes are broken down for each time point in Error! Reference source not found.. Many of the genes in this pathway encode ribosomal proteins either in the small or large subunit. However, there is little overlap between the ribosomal DEGs at these three time points. It is also interesting to note that while all the DEGs in this pathway are down-regulated for E14.5 and E16.5 time points, all the DEGs except for one are up-regulated at PND0. This result could indicate a defect in translation at the ribosome level and in protein synthesis, while the switch in the direction of fold-changes highlights the temporal aspect of regulation.

**Figure 7.**
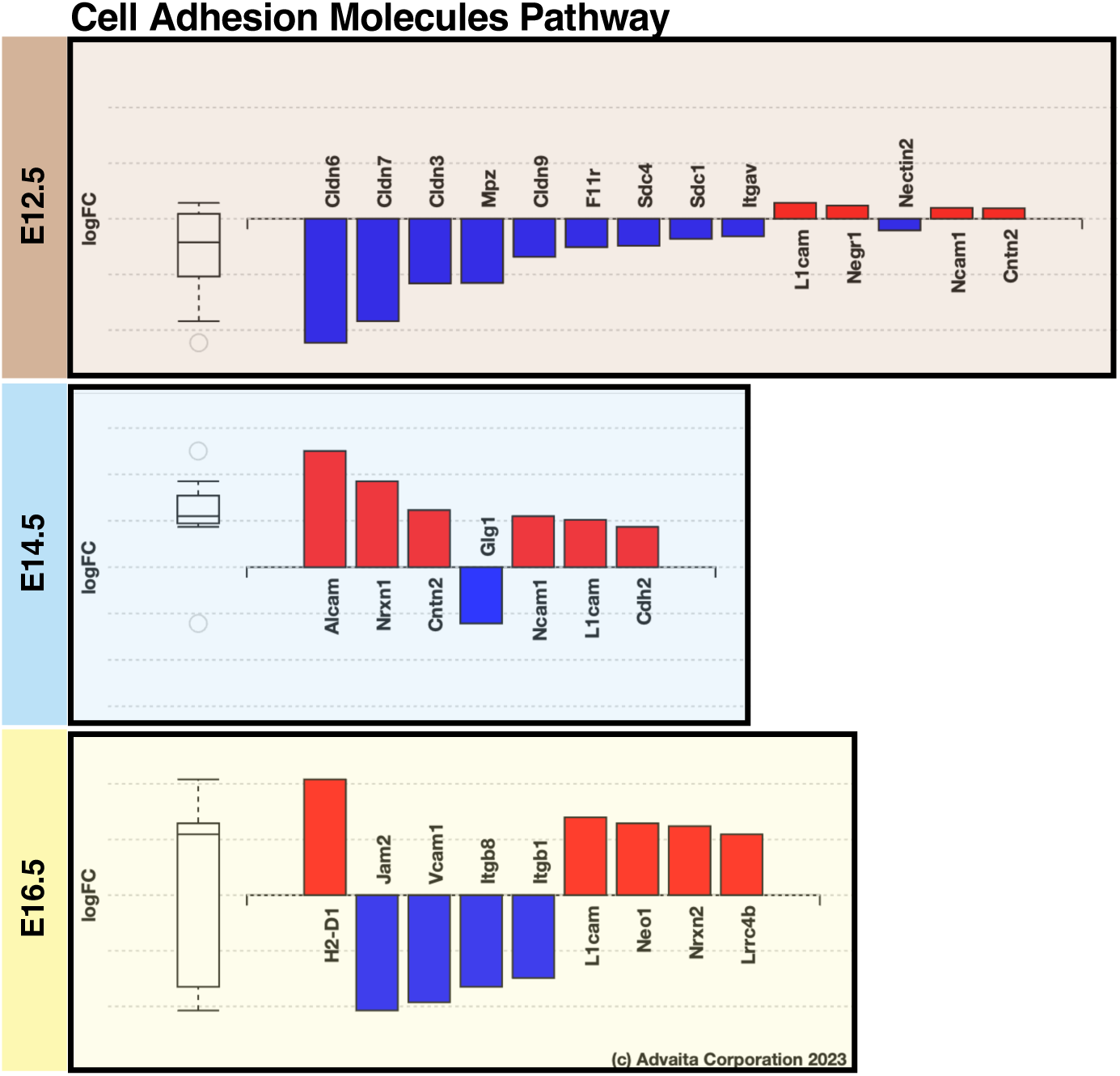
Bar plots of log fold-changes for DEGs in the **tight junction** (mmu:04530) and **cell adhesion molecules pathways** (mmu:04515) for time points where these pathways had a p-value ≤ 0.05.

Both *cholinergic* (mmu:04725) and *GABAergic synapse pathways* (mmu:04727) were significantly affected at the later E16.5 and PND0 time points (). For both, most genes are up-regulated at E16.5 and down-regulated at PND0, although the DEGs specific to the pathway differ between the time points. Gene expression changes in neurotransmitter signaling pathways could suggest defects in neurotransmission in SLOS, which will be discussed in the next section. Another related pathway, the *glutamatergic synapse* (mmu:04724), was shared between E12.5 and E16.5 time points (**Figure 10**), along with the *axon guidance* (mmu:04360) and *neuroactive ligand-receptor interaction pathways* (mmu:04080). Similar to the *cholinergic* and *GABAergic synapse pathways*, DEGs were found to be mainly up-regulated at the E16.5 time point. GABA is the primary inhibitory neurotransmitter, and glutamate is the main excitatory neurotransmitter in the mammalian CNS, suggesting that both inhibitory and excitatory neurotransmission were affected.

**Figure 8.**
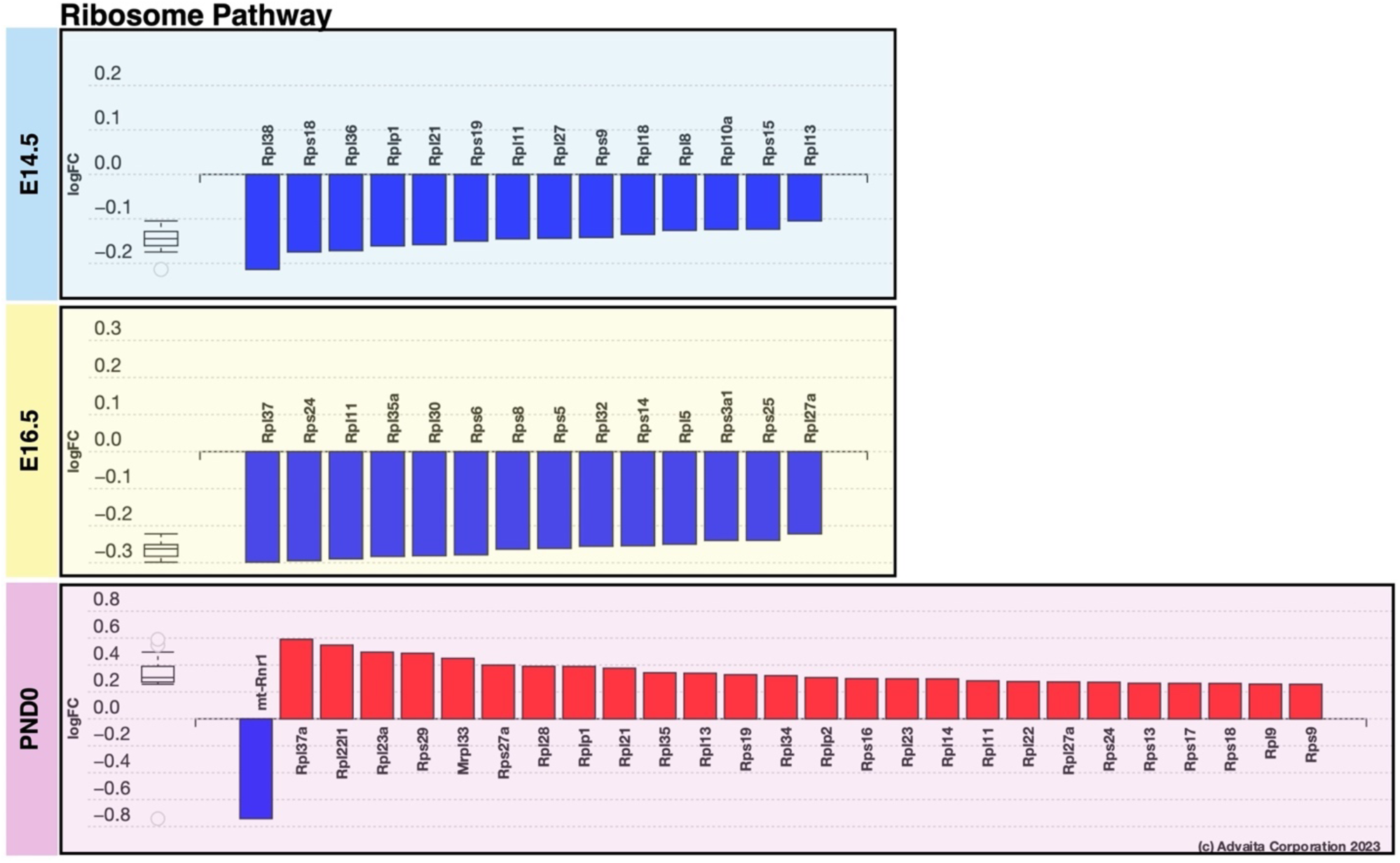
Bar plots of log fold-changes for DEGs in the **ribosome pathway** (mmu:03010) for time points where this pathway had a p-value ≤ 0.05.

**Figure 9.**
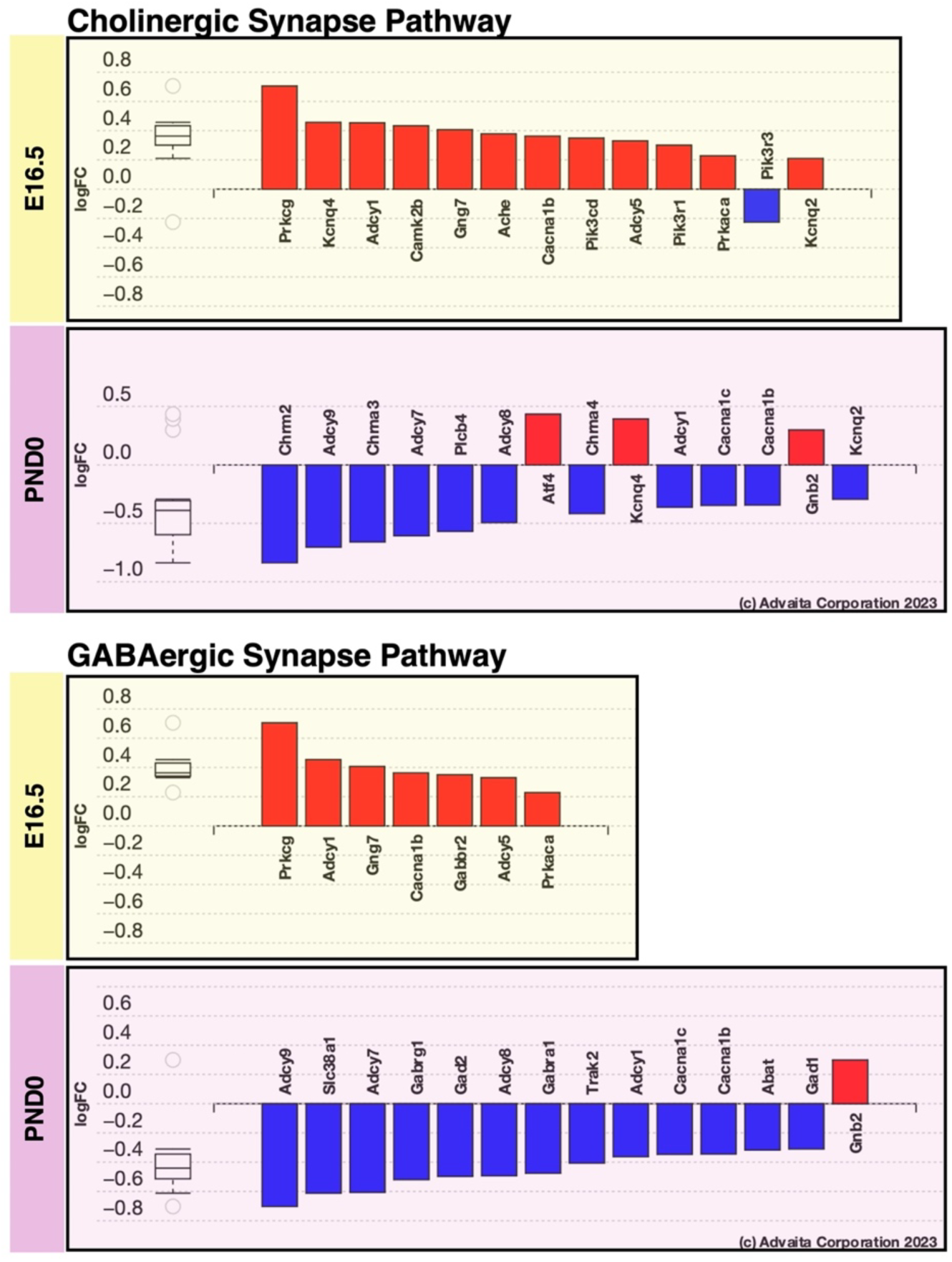
Bar plots of log fold-changes for DEGs in the **cholinergic** (mmu:04725) and **GABAergic synapse pathways** (mmu:04727) for time points where these pathways had a p-value ≤ 0.05.

**Figure 10.**
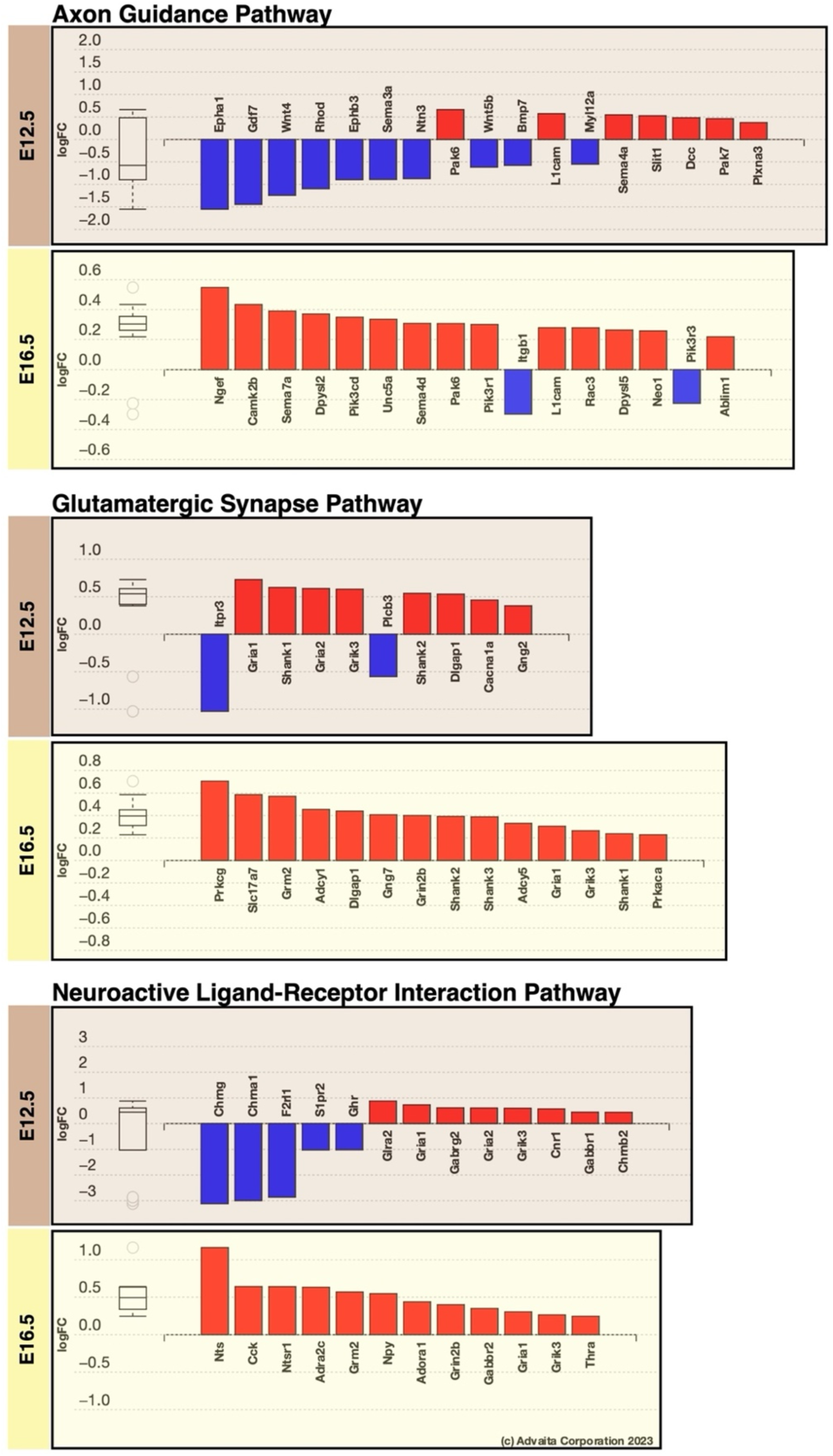
Bar plots of log fold-changes for DEGs in the **axon guidance** (mmu:04360), **glutamatergic synapse** (mmu:04724), and **neuroactive ligand-receptor interaction pathways** (mmu:04080) for time points where these pathways had a p-value ≤ 0.05.

### 3.4 Unique Pathways for Individual Time Points

In addition to the pathways commonly enriched at multiple time points, some are uniquely enriched at individual time points, which reflect the signaling pathways that are specific to the different stages of development. Thus, it would be reasonable to observe that amongst earlier time points, there were pathways more closely related to neurogenesis and early embryonic development of the CNS. In contrast, later time points had a greater emphasis on neuronal maturation and synaptic signaling.

### E12.5

Amongst the pathways significantly affected at E12.5 (Error! Reference source not found.**4**), three are related to embryogenesis, including *Hippo*, *Wnt*, and *TGF-β* pathways. All three are well-known, evolutionarily conserved signaling pathways crucial for mammalian embryonic development. DEGs for these three pathways are shown in an interaction network in **Figure 11**, where lines denote different types of interactions between genes and colors represent the direction of fold change. There is some overlap between the genes within these three related pathways. Most of the DEGs are down-regulated, which could indicate that inhibition of these pathways is underlying some of the structural and functional CNS abnormalities present in the SLOS phenotype.

**Figure 11.**
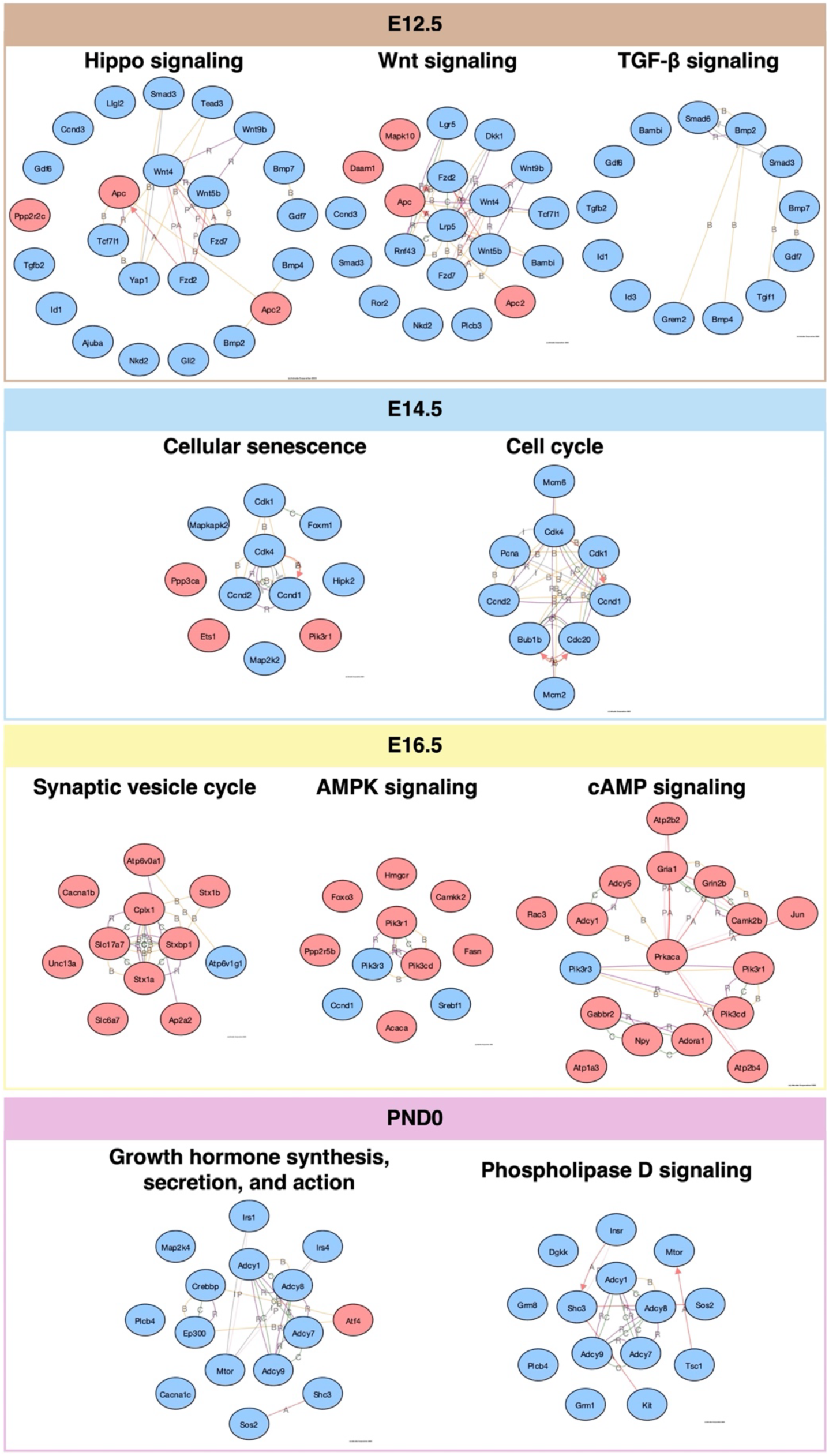
Interaction networks of DEGs in unique signaling pathways that are significantly affected at each time point (p-value ≤ 0.05). Up-regulated genes are shown in red, and down-regulated genes are shown in blue.

### E14.5

For E14.5 (mid-neurogenesis), two related pathways, *cell cycle* (mmu:04110) and *cellular senescence* (mmu:04218), were found to be among those significantly affected. Both pathways are related to cell cycle control, an important element of embryonic development and neuronal differentiation. Cyclin-dependent kinases (CDKs) are key regulatory enzymes in both pathways [46], and *CDK1* (the major CDK enzyme that controls cell cycle) and *CDK4* gene transcripts were both found to be down-regulated, as seen by the interaction network in **Figure 11**. This finding is interesting, considering the critical role of apoptosis in neurogenesis.

### E16.5

E16.5 had the largest number of unique pathways significantly affected among all the time points, with 31 pathways with a p-value ≤ 0.05. Among these, the *synaptic vesicle cycle* (mmu:04721) and two pathways related to *cAMP/AMPK signaling* (mmu:04024 and mmu:04152) were particularly interesting (**Figure 11**). Synaptic vesicles mediate the release of neurotransmitters into the synaptic cleft between synapses, and this affected pathway is related to the other synapse-associated pathways that were shared between several time points. Similar to those pathways, the DEGs for synaptic vesicle cycle were mostly up-regulated at E16.5. For cAMP/AMPK signaling pathways, there appears to be a link between these two pathways and their role in cellular homeostasis [47]. Dysregulation of cAMP/AMPK signaling could likely affect downstream physiological processes, including metabolism (various biosynthetic pathways), cell fate, and gene transcription.

### PND0

For PND0, we found many of the DEGs in significantly affected pathways to be down-regulated. In general, at this time point, there were more down-regulated genes than up-regulated ones, as seen in the asymmetrical volcano plot of DEGs (Error! Reference source not found.). Among the significantly affected pathways, *growth hormone, secretion, and action* (mmu:04935) and *phospholipase D signaling pathways* (mmu:04072) were of particular interest, as seen in **Figure 11**.

### 3.5 Loss of *Dhcr7* disrupts neurogenesis of ventral neural progenitors

It has been reported that *Dhcr7* is involved in modulating hedgehog and Wnt signaling [25, 48], which plays crucial roles in the development of GABAergic neurons and the ventral patterning of brain [49–51]. We recently elucidated that *Dhcr7* plays critical roles in neurogenesis in the dorsal cortices during development [39], but its role in ventral (GABAergic) neurogenesis has not been studied. Our transcriptome analyses have suggested that the lack of *Dhcr7* potentially affects the ventral neurogenesis during brain development. Thus, we examined whether *Dhcr7* is involved in ventral neurogenesis. To accomplish this, we used single embryo cultures of E11.5 *Dhcr7*^-/-^ (KO) or *Dhcr7*^+/+^ embryos. MGE/LGE were isolated from the ventral region for the single embryo cultures of neural progenitor cells. Three days after plating, the cells were immunostained for Ki67 and *β* III-tubulin (**Figure 12**). The results revealed that loss of *Dhcr7* led to a significant decrease in the proportion of Ki67^+^ precursors and a significant increase in the proportion of *β* III-tubulin^+^ neurons. Cleaved caspase 3(CC3) staining showed that loss of *Dhcr7* did not lead to increased apoptosis of ventral precursors in culture. These results indicated that loss of *Dhcr7* causes ventral neural progenitors to differentiate prematurely and decrease proliferation.

**Figure 12.**
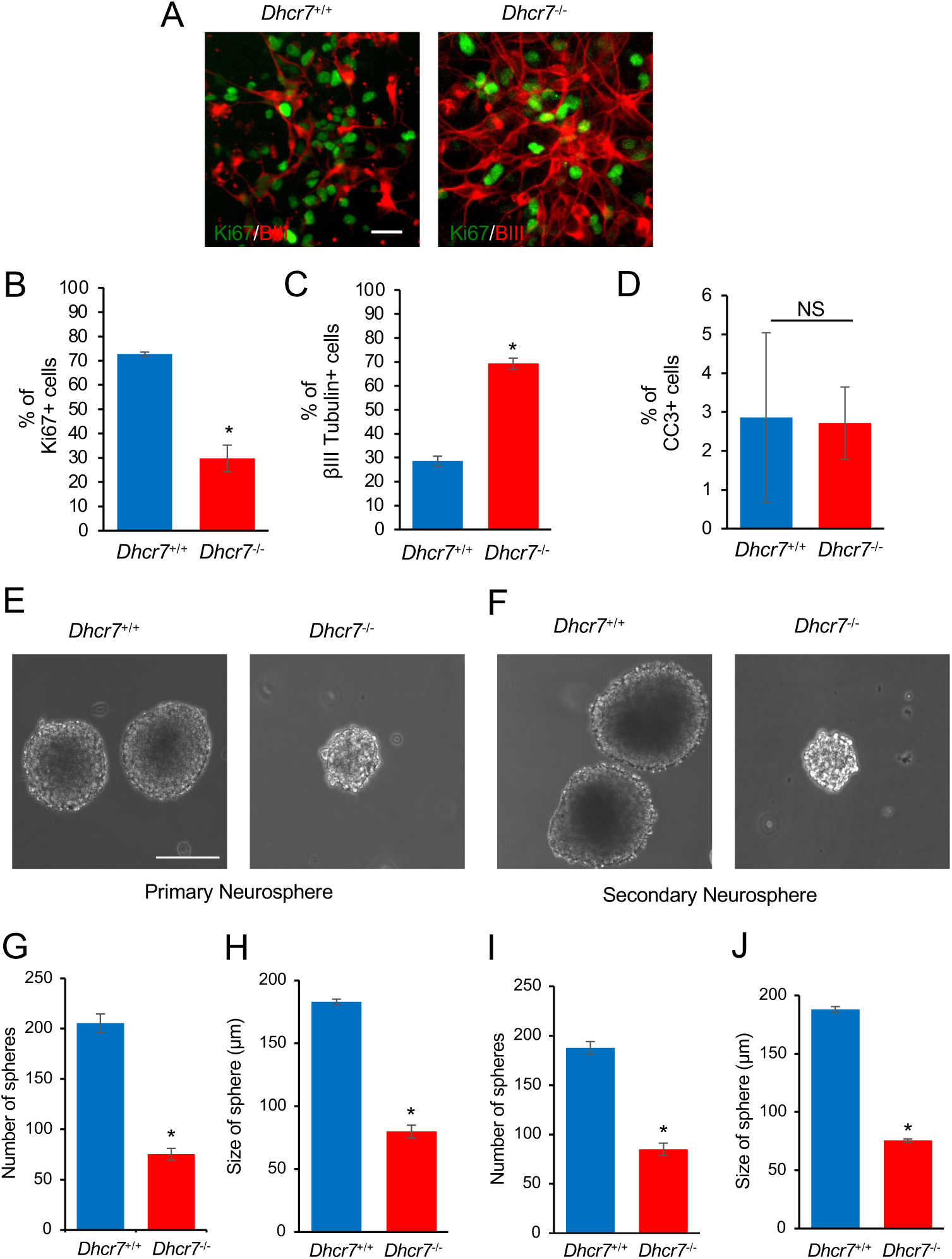
Loss of *Dhcr7* alleles causes decreased proliferation and increased neurogenesis in murine ventral cortical precursors. (A-D) E11.5 cortical precursors from single *Dhcr^+/+^* and *Dhcr7^-/-^* embryos were cultured 3 days and analyzed (A). Cells were immunostained for Ki67 (green) and βIII-tubulin (red) after 3 days and quantified for the proportions of Ki67+ (B), βIII-tubulin+ (C) and CC3+ cells (D). Scale bar = 50 μm. *, *p* < 0.001; n = 3 embryos per genotype. (E-J) E11.5 cortical precursor cells from single *Dhcr7^+/+^* or *Dhcr7^-/-^*embryos were cultured as primary neurosphere (E) and equal numbers of primary neurospheres were then passaged, generated secondary neurospheres (F). The number and diameter of primary and secondary neurospheres were quantified 7 days later (G-J). *, *p* < 0.001; n = 3 embryos per genotype. Scale Bar = 100 μm.

We then examined whether loss of *Dhcr7* potentially disrupts self-renewal of ventral neural progenitors by neurosphere assays, which evaluate whether progenitors can self-renew and repeatedly generate spheres. E11.5 ventral precursors from *Dhcr7*^-/-^ and *Dhcr7*^+/+^ embryos were cultured with FGF2 and EGF. The number and diameter of spheres were measured 7 days after plating. MGE/LGE from *Dhcr7*^-/-^ embryos generated significantly fewer numbers of and smaller neuropsheres than MGE/LGE from *Dhcr7*^+/+^ embryos. Following the formation of the primary neurpsheres, these spheres are triturated and re-plated at the equal density to form the secondary spheres. The results showed that the sphere size and numbers were approximately half the size of those from *Dhcr7*^+/+^ embryos. In parallel, we measured the sterol and oxysterol profiles of the neurospheres derived from *Dhcr7*^+/+^ and *Dhcr7*^-/-^ ventral neural precursors. Indeed, we observed accumulation of 7-DHC, 7-dehydrodesmostero (7-DHD), and 7-DHC-derived oxysterols (DHCEO, 4α-OH-7-DHC, 4β-OH-7-DHC, and 7-keto-8-DHC), as well as the deficiency of cholesterol, in *Dhcr7*^-/-^ neurospheres relative to *Dhcr7*^+/+^ (**Figure S3**), confirming the biochemical defects resulting from loss of *Dhcr7*, similar to those observed in *Dhcr7*^-/-^ mouse cortices [39]. Taken together, loss of *Dhcr7* significantly disturbs the proliferation and self-renewal of ventral progenitors.

## Discussion

Individuals with SLOS show a wide range of severity in phenotype across the many affected organ systems [9], and this applies to the neurological phenotype seen in the CNS. Cholesterol is particularly important for the brain because it is synthesized locally in the brain and is independent of an external source. Indeed, cholesterol plays a critical role in the developmentally essential Hedgehog and Wnt signaling pathways [15, 48, 52–57].

Cortical neurogenesis occurs over the span of several days, from approximately E11 to E17 [58]. *De novo* cholesterol synthesis in the mouse brain turns on around E10-E11, which coincides with the beginning of cortical neurogenesis [59]. The time points within our study span these neurogenesis stages. We and others have reported that loss of *DHCR7* leads to decreased proliferation and premature neurogenesis in both human and mouse NPCs [23, 25]. Francis et al. found that defective Wnt signaling may be responsible for such a phenotype, while our work suggested that activation of glucocorticoid receptor (GR) by an oxysterol metabolite may be the critical signaling pathway. However, these two findings are not mutually exclusive because activation of GR has been found to inhibit Wnt/β-catenin signaling [60, 61]. Wnt signaling is critical for basic developmental processes such as cell-fate specification, progenitor proliferation, and cell division [62]. At E12.5, *Wnt9b* (large fold-change), *Wnt4*, and *Wnt5b* are all down-regulated, as well as Wnt receptors, low-density lipoprotein receptor-related protein 5 (*Lrp5*) and Frizzled receptors (*Frz7* and *Frz2*). These transcriptomics results support the important role of Wnt signaling in the decreased proliferation and increased neurogenesis phenotype of SLOS NPCs.

*Hippo* and *TGF-β signaling pathways* are the other two significantly affected pathways at E12.5 that are critical for embryonic development. The Hippo signaling pathway controls organ size in embryonic development, which could be related to the commonly observed microcephaly phenotype in SLOS [9]. The signaling cascade consists of MST1/2 and LATS1/2 kinases and their cofactors [63]. In response to high cell density, activation of the pathway leads to cell apoptosis that prevents organ size overgrowth. At low cell density, when the pathway is inactivated, there are transcriptional factors that promote cell growth and proliferation. The TGF-β signaling pathway is involved in many different cellular functions, including proliferation, apoptosis, differentiation, and migration that are regulated by TGF-β family members. Most of the genes in Hippo and TGF-β signaling pathways are down-regulated, consistent with the decreased proliferation of *Dhcr7*-KO NPCs reported previously [23]. Previous work suggests that the Hippo pathway component LATS2 can inhibit SREBP [64], which regulates cholesterol biosynthesis. However, the mechanisms through which cholesterol deficiency affects the Hippo pathway remain to be elucidated. Furthermore, cholesterol has been found to suppress the TGF-β responsiveness by decreasing its binding to its receptor. Thus, how deficiency in cholesterol and/or accumulation of 7-DHC and its oxysterols affect TGF-β would be worth further investigation.

E14.5 is in the middle of cortical neurogenesis as NPCs differentiate into neurons that migrate to the outer layers of the neocortex while a proliferating pool of NPCs is also maintained [65, 66]. Programmed cell death, or apoptosis, plays a critical role in regulating neuronal precursors and pruning inefficient synapses [58]. Thus, it is not surprising to observe that cell cycle and cellular senescence were among the most significantly affected pathways at this time point. Indeed, previous work has shown that cholesterol is needed for CDK1 activation [67], which was downregulated in *Dhcr7*-KO brains at E14.5 (**Figure 11**).

It is important to note that GABAergic synapse, glutamatergic synapse, and cholinergic synapse pathways are significantly affected at later time points. Our previous work demonstrated premature neurogenesis of excitatory (glutamatergic) neurons using mouse cortical precursors [23]. In this work, we confirmed a similar phenotype of neurogenesis of GABAergic neurons from ventral neural precursors, i.e., premature differentiation and decreased proliferation (**Figure 12)**. These results indicate that the formation of all types of neurons may be affected similarly by the loss of *Dhcr7* function. Indeed, an impaired response of neurons to glutamate [68] and aberrant development of the serotonergic system [26] have also been reported previously.

At E16.5, the *axon guidance pathway* was found to be significantly upregulated. This is consistent with previous work by Jiang et al., which showed increased axon and dendrite formation in *Dhcr7*-KO mouse hippocampal neurons relative to WT [69].

Ribosome pathway was found to be significantly affected at E14.5, E16.5, and PND0, but mostly downregulated at E14.5 and E16.5 and upregulated at PND0. This is a novel pathway that has not been indicated in previous studies. A recent paper showed that reduced cholesterol levels block the proliferation of erythroid precursors by inhibiting ribosome biogenesis [70]. Although neural precursors are a different cell type from erythroid precursors, a similar mechanism could likely be in play for regulating the proliferation and differentiation of neural precursors. The role of cholesterol in regulating ribosome biogenesis in the brain and the significance of this pathway in neurodevelopment are worthy of further investigation in future studies.

Because SLOS patients are known to display autistic behavior [10, 11], we examined whether the DEGs observed in *Dhcr7*-KO mouse brains contain known genes involved in autism by comparing them with the SFARI autism gene database. Indeed, we found 69, 39, 75, and 114 autism-related genes at E12.5, E14.5, E16.5, and PND0, respectively (**Figure 13**). Among these genes, only *DHCR7* was shared by all time points. Genes shared by three groups include *FAT1*, *CACNA1B*, *KCNQ2*, *KCTD13*, *HERC2*, *STXBP1*, *LDLR*, and *TSPAN7*. Additional genes shared by at least two time points are shown in **Figure 13**.

**Figure 13.**
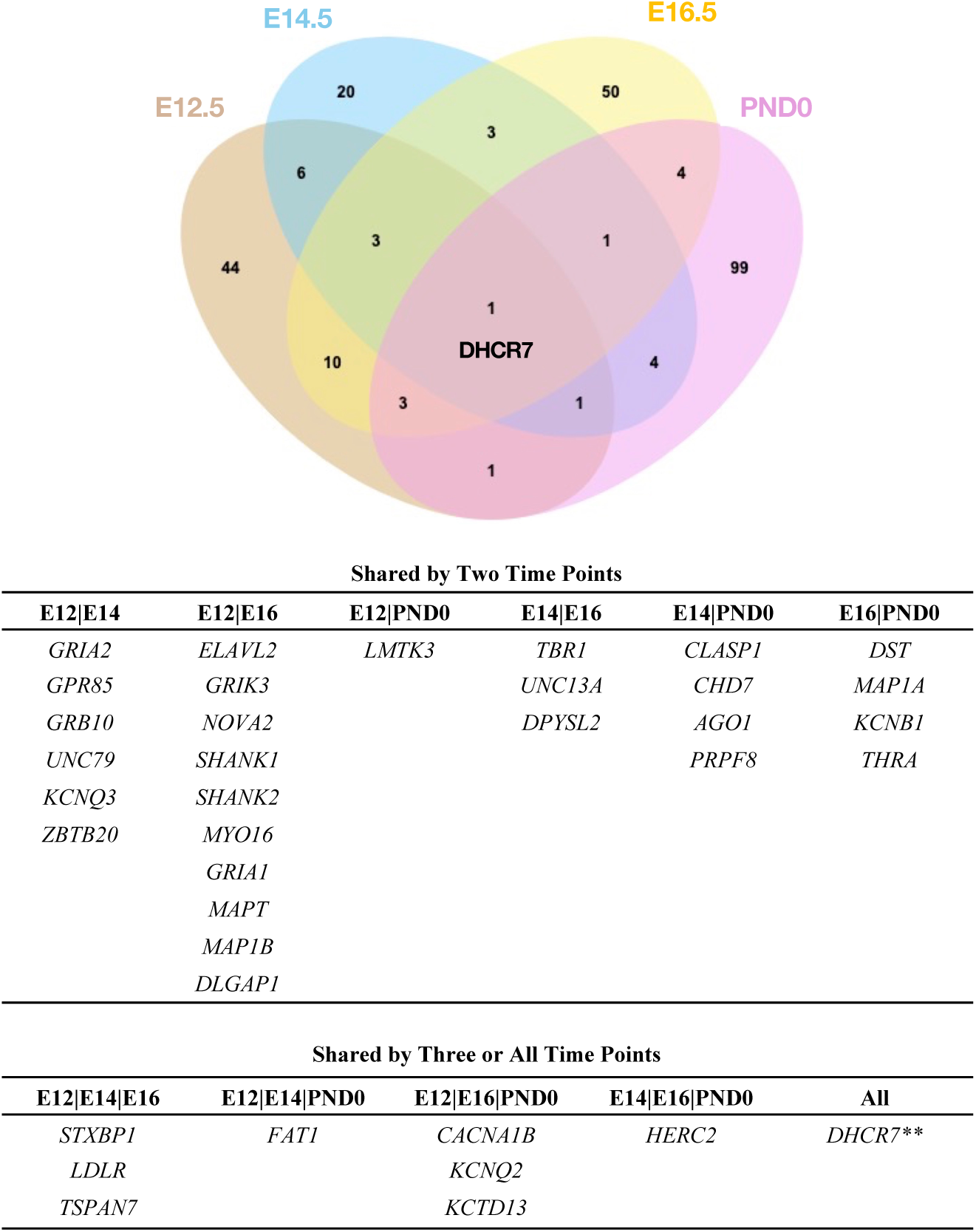
Venn diagram of DEGs shared with SFARI autism gene database at each time point.

## 4. Conclusion

In this work, we performed transcriptomic analysis of whole brains from *Dhcr7*-KO and WT mice at three embryonic time points and one postnatal time point. Comparative analysis of differentially expressed genes and pathway analysis shows the altered regulation of different physiological processes that could be related to the neurological phenotype in SLOS. Early time points suggest downregulation of genes and pathways involved in embryonic and neural development, such as Hippo, Wnt, and TGF-β, and later time points suggest universal defective neuronal synapse pathways. In addition, the ribosome was found to be another novel pathway affected by deficient cholesterol biosynthesis at three time points. *In vitro* neurogenesis experiments using mouse ventral neuronal precursors confirmed that loss of *Dhcr7* led to decreased proliferation and premature neurogenesis, consistent with the transcriptomic changes. Thus, disruption of cholesterol synthesis through genetic mutation of *Dhcr7* has deleterious and concerted effects beyond sterol and lipid homeostasis.

## Author Contributions

LX conceived the study. AL and LX designed the study. AL and HT collected samples. AL performed the experiments and data analysis related to transcriptomics. HT carried out the experiments related to the *in vitro* neurogenesis experiments. AL, HT, and LX wrote the manuscript.

## Acknowledgements

Funding: This work was supported by grants from the National Institutes of Health: R01 HD092659 (L.X.), Pharmacological Sciences Training Program (T32 GM007750), the NIEHS funded EDGE Center (P30ES007033), and Institute of Translational Health Sciences TL1 Program (TL1 TR002318) from the National Center for Advancing Translational Sciences. The content is solely the responsibility of the authors and does not necessarily represent the official views of the National Institutes of Health.

We would like to sincerely thank James W. MacDonald and Theo K. Bammler for their knowledge and assistance with iPathway Guide through the University of Washington EDGE Center.

