## Supplemental Figures S1 and S2 for "Temporal transcriptomic changes during neurodevelopment in a mouse model of Smith-Lemli-Opitz syndrome"

**Table of Content**

**Supplementary Table S1** List of differentially expressed genes for each time point with an adjusted p-value (FDR) ≤ 0.05. See separate Excel file.

**Supplementary Table S2** List of enriched KEGG pathways for each time point with a p-value ≤ 0.05. See separate Excel file.

**Supplementary Table S3** List of enriched GO Biological Process terms for each time point with an adjusted p-value (FDR) ≤ 0.05. See separate Excel file.

**Figure S1.** PCA bi-plots for **A.** all groups combined and **B.** individual time points.

**Figure S2.** Top 10 enriched GO terms using iPathwayGuide.

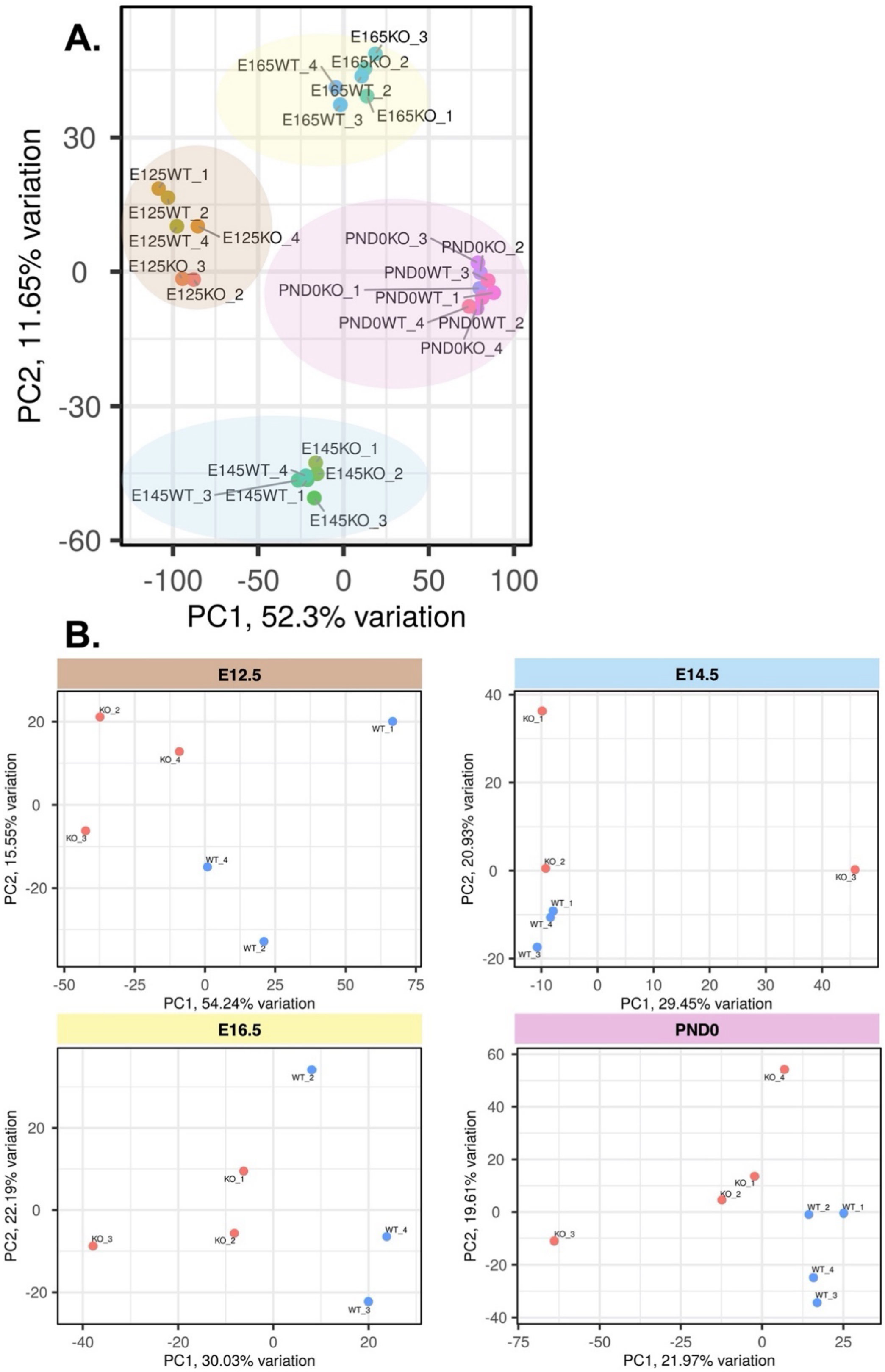


**Supplementary Figure S1.** PCA bi-plots for **A.** all groups combined and **B.** individual time points.
Created by iDEP1.1.

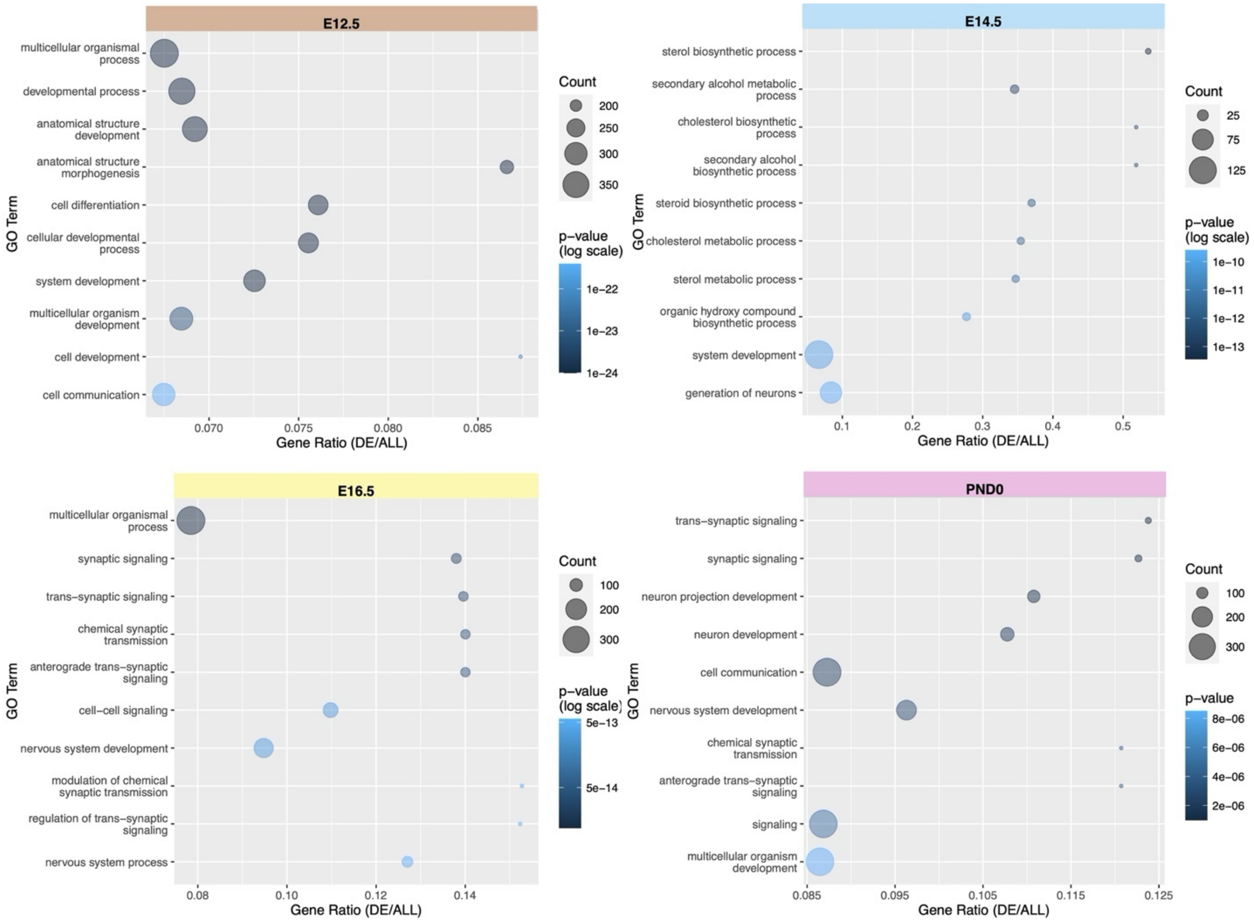


**Supplementary Figure S2.** Top 10 enriched GO terms using iPathwayGuide. Bubble plots are ordered by p-value and display pathway name and gene ratio, where the size of the bubble designates the number of differentially expressed genes and the color designates p-value. Created by R/ggplot2.
